# Normative neural and physiological correlates of parent-child observational threat extinction

**DOI:** 10.1101/2025.08.11.669679

**Authors:** Sara A. Heyn, Ryan J. Herringa

## Abstract

The dyadic relationship between a parent and child is a critical facilitator of social learning, particularly in the management of fear. Vicarious extinction, the process of learning by observing a parent extinguish learned threat responses, holds critical translational potential in understanding adolescent psychopathology involving aberrant threat responses. However, normative biological profiles of vicarious extinction have yet to be uncovered. Here, a community sample of 97 parent-child dyads, enriched for trauma-exposure, completed a validated vicarious extinction paradigm. Youth completed all phases during functional magnetic resonance imaging, while caregivers simultaneously completed the behavioral task. Linear models interrogating normative behavioral, physiological, and neural correlates of acquisition and direct and vicarious extinction, controlled for trauma and symptom severity. The degree of parent-child autonomic synchrony was used to predict the strength of extinction learning. Youth behavioral and skin conductance response markers suggest successful threat acquisition and direct and vicarious extinction. Autonomic arousal during active vicarious extinction learning processes was significantly associated with the strength of parent-child synchrony. Neural activation analyses reveal patterns of early encoding of threat and safety discrimination that were reactivated during later threat reinstatement. Finally, results corroborate and expand known models of direct and vicarious extinction, respectively involving the ventromedial and ventrolateral prefrontal cortex. Altogether, this data confirms that youth can directly and vicariously modify learned threat associations with distinct neurobiological profiles. This delineation of vicarious extinction neural circuitry within a normative population is pivotal for understanding how these processes may be altered by the environment and/or psychopathology during adolescence.

## I. INTRODUCTION

Within the framework of normative and atypical adolescent development, the importance of the parent-child dyadic relationship cannot be understated. There are many documented pathways for intergenerational learning and stress transmission[1], including through genetics[2], prenatal programming[3], or vicarious threat learning (the process of learning to discriminate threat and safety via observation)[4,5]. Social learning theory has extensively theorized that children can adaptively attenuate their own learned threat responses simply by observing how their parent responds to that threat[6]. Disruption of this process may underlie risk and resilience of psychopathology, wherein youth that are unable to adapt and diminish a learned threat response via parental observation may be susceptible to clinical dysfunction. This may mirror known associations between aberrant threat extinction and anxiety and posttraumatic stress disorder (PTSD) in adults[7,8]. Despite strong translational potential, the study of vicarious threat extinction in childhood is in its infancy, notably lacking a foundational assessment of how this transmission occurs in normative adolescent populations.

Evaluations of threat and safety learning processes typically harness the simplicity of Pavlovian conditioning paradigms[9] that can rapidly generate threat associations across humans and animals[10]. In brief, a neutral stimulus (conditioned stimulus, CS) is paired with an aversive, unconditioned stimulus (US; e.g., loud noise, electrodermal shock) to induce a threat response (e.g., freezing behaviors, increased skin conductance response, US expectancy). Direct extinction of this association is learned through repeated exposure to the CS without the US. Finally, reinstatement acts as a robust assessment of extinction learning stability and ongoing vulnerability to threat exposure[11].

Recent reviews of the threat extinction literature present the prevailing neurobiological model of threat extinction (i.e., the fear extinction network, FEN)[10,12]. The FEN, generally responsible for threat detection and response, typically includes the amygdala, dorsal anterior cingulate cortex (dACC), hippocampus, and ventromedial prefrontal cortex (vmPFC)[13], with suspected related nodes in the insula, cerebellum, thalamus, and periaqueductal gray[14]. The strength of these inter- and intra-network connections undergo critical developmental changes during adolescence.[10]

Notably, these well-established models have yet to incorporate mechanisms of vicarious threat extinction, particularly within the adolescent period when parent-child dyadic relationships remain developmentally salient. In fact, from a developmental perspective, early experiences of threat and safety learning are often based upon vicarious or observational experiences rather than direct learning events[15], may represent a critical and adaptive survival mechanism, and have been suggested to be just as effective as direct experiences in adults[16,17]. Young children don’t need to touch a fire to know that its hot, adolescents don’t need to get bit by a snake to be afraid of seeing a snake while on a hike. Vicarious threat acquisition in parent-child dyads can be physiologically quantified[5] and deviations may be relevant to anxiety and related psychopathology[18]. In contrast, the hallmark re-experiencing symptoms of PTSD, a clinical manifestation of dysregulation following trauma[19], suggests alternate or additional impairments in threat extinction. It may be that the inability to vicariously extinguish these associations (i.e., persistent fear despite repeated exposures to their parent modeling composure when recounting trauma) may be a more specific and potent predictor of posttraumatic stress disorder (PTSD) development and persistence during adolescence.

Vicarious extinction learning, which requires the integration of safety signals modeled by others during the extinction process, presumably relies both upon the FEN and parallel networks that facilitate mentalization and emotion attribution[20]. Of note, the emotion attribution (EA) network is hypothesized to recognize and infer causal attribution of others’ emotional states. The EA is thought to integrate signals from the anterior temporal cortex (aTC), superior temporal sulcus (STS), and the dorsomedial (dm-) and ventrolateral (vl)PFC. In fact, a meta-analysis of vicarious learning during reward processing paradigms in adults suggests preferential engagement of regions with the EA network. So, while vicarious extinction learning may reflect a unique, interdependent process involving both FEN and EA network nodes, normative studies in adolescents must substantiate these hypotheses.

We have previously developed and validated a three-day behavioral paradigm of direct and observational threat extinction in youth with and without trauma exposure and their caregiver[4]. This behavioral data supports the use of this paradigm to quantify parent-child vicarious extinction and is tolerable within trauma-exposed youth with PTSD and suggests possible physiological mechanisms of extinction learning transmission within parent-child autonomic synchrony. Dyadic synchrony is the moment-to-moment temporal concordance of micro-level behavioral, neural, or physiological signaling, and has been conceptually proposed as a unique marker of intergenerational transmission[21]. In particular, coordinated autonomic activity may indicate reciprocal, co-created arousal[22], making it an optimal target for quantifying threat extinction processes that are reflective of arousal states.

To our knowledge, this is the first comprehensive assessment of functional mechanisms of direct versus vicarious threat extinction learning across behavioral, autonomic, neural, and parent-child synchrony within a community sample of parent-youth dyads. We hypothesize to see recruitment of nodes within the FEN (amygdala, vmPFC) during direct extinction, recruitment of nodes within the EAN (dmPFC, vlPFC) during vicarious extinction, and associations between vicarious extinction learning and dyadic autonomic synchrony. The characterization of common biological profiles of vicarious extinction will provide a critical baseline for examining profile modifications that may aid in predicting diagnosis of fear-related disorders as well as potential therapeutic responses in trauma-exposed youth.

## II. METHODS

### Participant recruitment and behavioral assessments

A community sample of 97 parent-child dyads, youth ages 10-14 years (inclusive), were recruited to complete a three-day observational threat extinction learning paradigm. Recruitment methods including community and social media flyers, school newsletters, pediatric health clinics, university listservs, and messaging through electronic medical records to potentially eligible families.

This study recruited both healthy, typically developing youth as well as youth with trauma exposure with and without posttraumatic stress disorder (PTSD). Inclusion criteria for the full sample included: (1) both parent and youth willing to participate and provide consent/assent in English; (2) youth and caregiver with visual acuity to read text on a computer monitor; (3) youth 10-14 years of age, inclusive; (4) youth is able to lie still for up to two hours, and (5) the youth is willing and able to safely complete and MRI scan. Exclusion criteria for the full sample included: (1) a medical condition that would prevent study participation; (2) caregiver with a current or history of perpetrating abuse against the youth participant; (3) current pregnancy; (4) youth with ongoing trauma, active substance abuse, imminent suicidality, current or past bipolar, psychotic, neurological (e.g. epilepsy) or neurodevelopmental (e.g. autism spectrum) disorders; (5) yellow-blue color blindness; (6) traumatic brain injury with ongoing symptoms; (7) medications that may interact with study activities (e.g. direct impacts on the sympathetic nervous system). This project was approved by the University of Wisconsin-Madison Health Sciences Institutional Review Board, all study procedures were conducted in accordance with the approved protocol, and all parents and youth provided written informed consent and assent, respectively. A total of 87 parent-child dyads were eligible, interested, and completed at least one day of the full paradigm. Two dyads only completed day one and three additional dyads only completed days one and two, resulting in 82 dyads completing the full paradigm.

Youth and parents underwent a full clinical diagnostic and trauma assessment protocol, including web-based clinician administration of the Kiddie Schedule for Affective Disorders for the DSM-V (KSADS-CA)[23] and Yale-Vermont Adversity in Childhood Scale (Y-VACS)[24]. Current mental health symptoms were assessed using the Mood and Feelings Questionnaire (MFQ; assessment of depression symptoms)[25], Screen for Child Anxiety-Related Disorders (SCARED; assessment of anxiety symptoms)[26], and the UCLA Posttraumatic Stress Disorder Reaction Index for the DSM-V (PTSD-RI; assessment of PTSD symptoms)[27]. Due to lack of follow-up, a small subset of subjects were missing PTSD-RI scores (n=6).

### Observational threat extinction paradigm

Parents and youth simultaneously completed an adapted three-day observational extinction learning paradigm (Figure 1)[4,28]. Youth completed the entire paradigm while undergoing functional magnetic resonance imaging while parents completed the behavioral paradigm. During each phase of the paradigm, youth and parent wore a respiration belt, had two electrodes on the collarbone and opposite hip, Ag/AgCl SCR electrodes (EL509, isotonic recording GEL 101) attached to the pointer and middle finger of the left hand, and two electrodes on the pointer and middle finger of the right hand delivered electrodermal shocks as the unconditioned stimulus (US). Both participants underwent shock calibration procedures to choose an intensity that was annoying but not painful and the chosen intensity was preserved throughout the paradigm.

**Figure 1.**
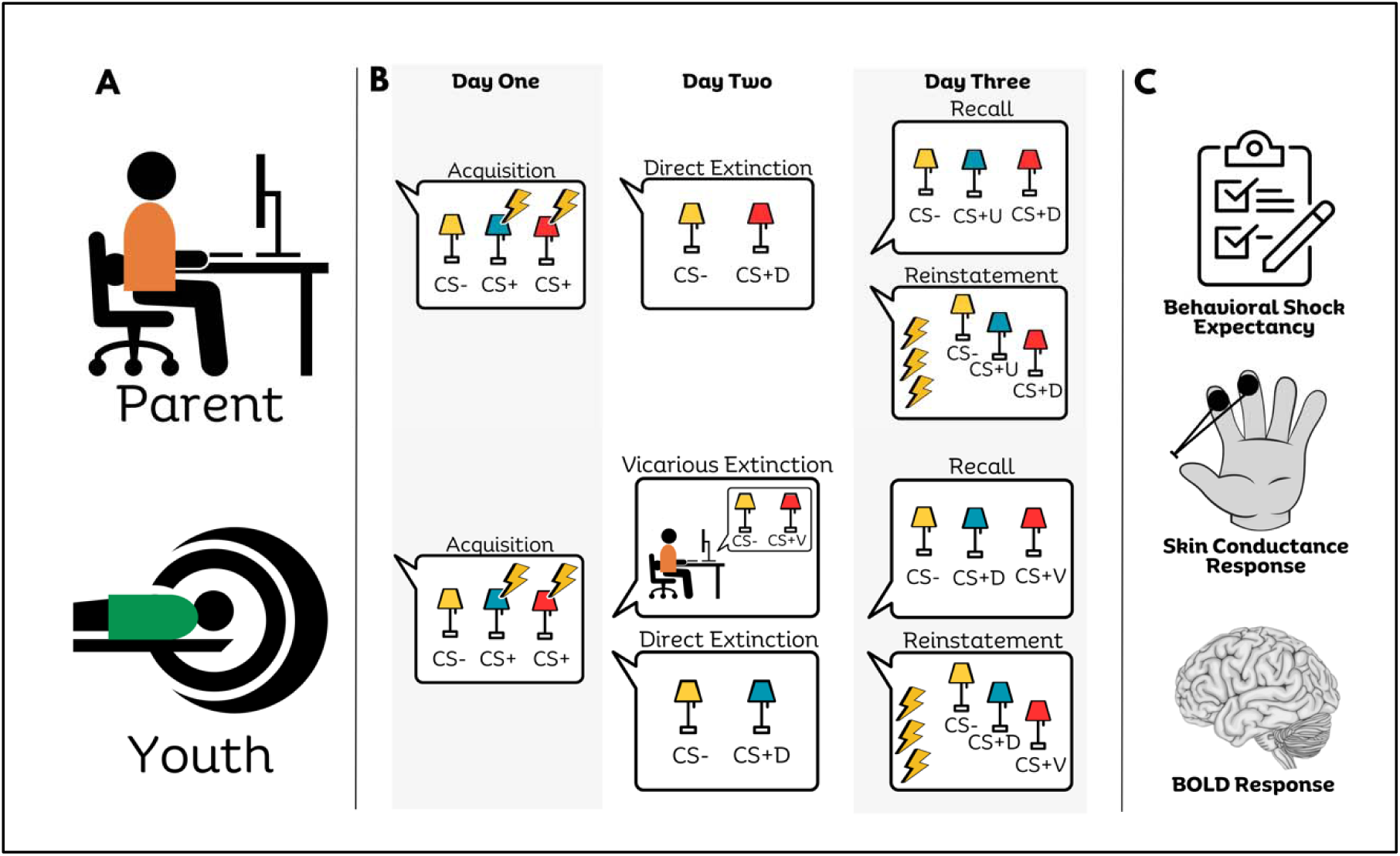
Task Design and Primary Outcome Variables. (A) This three-day paradigm assessing vicarious extinction learning in youth was completed by youth in the scanner and a caregiver. (B) A schematic depicting the conditioned stimuli (yellow lamp, CS-; blue lamp, CS+Dir; red lamp, CS+Vic) paired with the unconditioned stimuli (US; electrodermal shock) during acquisition (50% reinforcement rate). Both youth and caregivers’ complete direct extinction (CS+Dir presentation with no US-pairing) while youth additionally complete vicarious extinction (viewing of a video of their caregiver completing direct extinction). All participants completed extinction recall (no US-pairings) and reinstatement (unpaired US presentation prior to task start) on day three. (C) Primary outcome variables included self-report of US expectancy, skin conductance response (SCR), and BOLD activation.

On Day 1, participants completed a brief habituation phase, where three colored lamps (yellow, red, and blue) were presented without US-pairing, followed by threat acquisition. Acquisition consisted of two counterbalanced blocks of CS-US pairings with two different lights (red, blue) and the unpaired CS- (yellow). There was a total of 8 CS+ trials for each CS+, presented with a 50% reinforcement rate. Each CS trial (6s) was immediately preceded by an unlit lamp (3s) and followed by a jittered inter-trial interval (9-14s). During acquisition, the lamps were presented in one context (i.e., on a computer desk), while all proceeding phases were presented in a separate context (i.e., on a bookshelf). On Day 2, parent participants completed direct extinction learning, during which 6 trials of one CS+D (CS+Direct; red) and 4 trials of the CS- (yellow) were presented with no US-pairing. Parents underwent direct extinction training while being video recorded. Similarly, youth participants completed direct extinction learning during which 6 trials of one CS+ (CS+Direct; blue) and 4 trials of the CS- (yellow) were presented with no US-pairing. Youth also completed vicarious extinction learning, during which youth participants watched the video recording of their parent undergoing direct extinction (CS+Vicarious; red). The order of direct and vicarious extinction was randomly counterbalanced. On Day 3, both youth and parent participants completed extinction recall and threat reinstatement. During recall, 8 trials of all three lamps (red, blue, yellow) were presented without US-pairing. During threat reinstatement, prior to the start of the task, participants received a series of three unpaired shocks while viewing a black screen followed by an additional 8 trials of all three lamps (red, blue, yellow) without US-pairing.

Before and after each phase, in addition to attentional checks, participants completed an adapted behavioral shock expectancy questionnaire [29]. Here, they estimated, “On a scale from 1 (not at all) to 5 (very much), how much did you expect a shock for the [first, last] trial of [blue, yellow, red]?”. To assess threat acquisition, extinction, and recall, expectancy scores were then analyzed with a series of linear mixed effect models: (1) an omnibus model across all six phases (habituation, acquisition, direct extinction, vicarious extinction, extinction recall, and reinstatement) estimating a phase by stimulus type (CS+Dir, CS+Vic, CS-) by order (first trial, last trial) interaction, and (2) separate models within each phase estimating a stimulus type (CS+Dir, CS+Vic, CS-) by trial number interaction. All models included youth age, youth sex, cumulative adversity composite score, and PTSD symptom severity, as well as subject as a random effect. Significant interactions were decomposed with subsequent linear mixed effects models, where appropriate. Given the specific sample enrichment for PTSD and high collinearity between PTSD symptoms and anxiety [*r*(76)=0.59, *p*<0.0001] and depression [*r*(77)=0.49, *p*<0.0001] symptoms[30], sensitivity analyses specifically controlling for these alternate symptom domains were not assessed. For brevity, we report only models from youth participants.

### Autonomic psychophysiology

#### Skin conductance recording and processing

For both youth and parent, skin conductance levels (SCL) were continuously collected (MP160 recording system, Biopac Systems Inc., Goleta, CA) directly into BIOPAC AcqKnowledge software at 1000Hz. According to previous recommendations [31], for each phase, SCL underwent (1) bi-directional first-order Butterworth filtering using a high-pass of 0.03Hz and low-pass of 5Hz, (2) downsampling to 50Hz for computational efficiency, and (3) Z-scored. Consistent with previous paradigms [4,32], skin conductance responses (SCRs) were obtained via subtracting the mean SCL during the immediately preceding unlit lamp (3s) from the peak SCL during the 6s stimulus presentation. Finally, SCRs were square-root transformed.

To assess threat acquisition, extinction, and recall, SCRs were then analyzed with a series of linear mixed effect models: (1) an omnibus model across all six phases (habituation, acquisition, direct extinction, vicarious extinction, extinction recall, and reinstatement) estimating a phase by stimulus type (CS+Dir, CS+Vic, CS-) by trial number interaction, and (2) separate models within each phase estimating a stimulus type (CS+Dir, CS+Vic, CS-) by trial number interaction for post-hoc decomposition of within-phase dynamics. All models included youth age, youth sex, cumulative adversity composite score, and PTSD symptom severity, as well as subject as a random effect. Significant interactions were decomposed with subsequent linear mixed effects models. For brevity, we report only models from youth participants.

#### Parent-child extinction training physiological synchrony

Parent-child physiological autonomic synchrony analyses were performed for the extinction sessions (vicarious extinction while youth view a video of their parent completing direct extinction training). Consistent with previous studies [32], raw SCLs underwent filtering using a 0.05-1Hz passband and downsampled to 50Hz for computational efficiency. Finally, the time series were Z-scored to ensure comparability between youth and parent. To quantify the time-dependent coupling of parent and child physiological signal during extinction training, a bivariate extension of recurrences quantification analyses, cross-recurrence quantification analysis (CRQA), was performed. Analyses were completed in R using the *crqa* package.

In summary, CRQA yields a cross-recurrence plot of phasic skin conductance signal from each parent and child that represents the state of recurrence between the individual time series (i.e., within a given radius). The optimal parameters (embedding and delay dimensions, radius) were individually determined for each pair of signals to yield an average recurrence rate between 2-4% [33], which represents the percentage of points in the cross-recurrence plot that form recurrence. Once the cross-recurrence plots were created, a series of synchrony metrics were extracted to mirror previous studies [32,33]. Determinism (DET) and laminarity (LAM) exemplify the percentage of recurrence points that form diagonal and vertical lines, respectively. The maxL metric represents the length of the longest diagonal in the cross-recurrence plot (maxL). Finally, entropy (ENTR) typifies entropy of the diagonal line length distribution within the cross-recurrence plots [34]. The maxL metric was highly uncorrelated with the other three metrics and was ultimately excluded. Data reduction to a signal synchrony component for each dyad was performed using a principal component analysis (PCA) with varimax-rotation. The first component of this PCA was used to separately predict behavioral expectancy and skin conductance response during the final two trials of vicarious extinction and first two trials of extinction recall for the CS- and CS+Vicarious, controlling for youth age, youth sex, cumulative adversity composite score, PTSD symptom severity, subject as a random effect.

Of the participating 82 parent-child dyads with completed paradigms, 7 dyads were excluded due to late starts in physiological data acquisition, 2 dyads were excluded due to being SCR non-responders, and 1 dyad was excluded due to being unable to properly optimize parameters to obtain an average recurrence rate between 2 and 4%. Therefore, a total of 73 parent-child dyads were included in SCR and 72 dyads in synchrony analyses.

### Functional magnetic resonance imaging

#### Acquisition Parameters

All MRI data were acquired using a Discovery MR750 3.0T scanner (GE Healthcare, Chicago, IL) equipped with a 32-channel head coil. At the beginning of each day of scanning, a high-resolution T1-weighted axial 3D MPRAGE scan (TR=2.5s, TE=0.3ms, matrix size:256x256, FOV=100mm, slice thickness=1mm, flip angle =8°) was performed. During each task phase, blood oxygen level-dependent (BOLD) signals were captured using a T2-weighted multiband gradient echo planar imaging (EPI) sequence (TR=1.6s, TE=0.025s, matrix size: 90x90, FOV=100mm, slice thickness=2.4mm, flip angle=65°, multiband acceleration factor=3).

#### fMRI first-level processing

Initial preprocessing of the functional images was performed using fMRIPrep 23.2.0[35] (RRID:SCR_016216), which is based on Nipype 1.8.6 [36](RRID:SCR_002502). Images from all three days were longitudinally registered and processed using default fMRIPrep parameters and a fieldmap-less distortion correction approach (susceptibility distortion correction; SDC). Full details are included in Supplemental Methods. Quality control methods utilized MRIQC [37] and fMRIPrep html outputs. We followed previously published guidelines for initial quality review [38] in tandem with the removal of any phase with a mean framewise displacement (FD) > 3SD above the mean. Altogether, QC methods resulted in the removal of 7 individual phases across three subjects (habituation, n=1; acquisition, n=1; direct extinction, n=2; vicarious extinction, n=1; reinstatement, n=2).

Next, all phases underwent the following additional first-level processing procedures: (1) smoothing using a 4.0mm FWHM kernel, (2) scaling, (3) nuisance regression with AFNI’s 3dDeconvolve, including 128sec high-pass filter, 6 motion regressors convolved with a gamma HDR function, and slice timing corrections [39]. Of note, we did not include global signal response (GSR)[40] or FD (due to QC procedures) regressors. Finally, to estimate individual betas for each trial of each phase (including the US, although US-specific effects are outside the scope of this paper) and account for signal drift and biorhythms, we fit a first-order autoregressive (AR) model using the REML generalized least squares (GLSQ) regression method in AFNI’s 3dREMLfit. Final voxel resolution was 2mm x 2mm x 2mm.

This methodology resulted in an individual beta map for each trial of each conditioned stimulus and each preceding context stimulus within each phase. Next, a beta contrast map removing the variance of the preceding context stimulus from each trial of each conditioned stimulus was created. Finally, an average of the first two and last two conditioned stimuli (CS+Dir, CS+Vic, and CS-) for each phase was created and input in final analytical models. To comprehensively assess neural mechanisms of threat acquisition, extinction, recall, and reinstatement, we conducted both whole-brain and region-of-interest (ROI) analyses.

Whole-brain analyses employed linear mixed-effect modeling within AFNI’s 3dLMEr using a gray matter mask, including an omnibus model estimating a phase (habituation, acquisition, direct extinction, vicarious extinction, recall, reinstatement) by stimulus type (CS+Dir, CS+Vic, CS-) by phase order (early trials, late trials) interaction in addition to individual models within each phase, estimating a stimulus type (CS+Dir, CS+Vic, CS-) by phase order (early trials, late trials) interaction. Each model included youth age, youth sex, total adversity composite score, and PTSD symptom severity, with subject as a random effect. A conservative approach to multiple comparison correction via cluster simulations was employed using AFNI’s 3dClustSim, which computes a cluster-size threshold for a given voxelwise threshold of (p_uncorr_<0.005) for each phase. We report significant clusters that survive p_cor_<0.05 according to the following cluster thresholds (NN=3): acquisition, k=80; direct extinction, k=80; vicarious extinction, k=86; extinction recall, k=73; and reinstatement, k=73. Finally, region-of-interest analyses assessed identical models described above within 66 pre-identified ROIs hypothesized to be associated with threat extinction networks. Using the Schaefer 400 atlas [41], average betas from each stimulus, trial order, and phase for each participant was extracted from the following ROIs: bilateral dorsomedial prefrontal cortex (dmPFC; n=17)[42], ventromedial PFC (vmPFC, n=18), ventrolateral PFC (vlPFC, n=10), dorsal anterior cingulate cortex (dACC, n=7)[43]. Similarly, average betas were extracted from subcortical regions using the Tian IV atlas [44], including the amygdala (n=4) and hippocampus (n=10). Further detail regarding ROI identification can be found in Supplemental Methods. To correct for multiple comparisons, only ROIs surviving false discovery rate (FDR) correction are reported.

## III. RESULTS

### Participant Characteristics and Behavioral Task Validation

Parent-child dyad socio-demographics are presented in Table 1. While this paradigm has been validated in a previous study[4], analyses of behavioral shock expectancy provide validation for successful threat acquisition and extinction for this paradigm in the MR scanner. Decomposition of a significant phase by stimulus by order interaction [F(8,1789)=19.67,*p*<0.001] in stimulus by order interactions within each phase show increased shock expectancy for both CS+ and decreased shock expectancy for the CS- by the final trial of threat acquisition [F(2,362)=39.62,*p*<0.001] and decreasing shock expectancy for the CS+ by the last trial of direct extinction [F(2,228)=15.92,*p*<0.001] . During vicarious extinction, we see only a significant effect of stimulus for the CS+V [F(2,255)=23.5,*p*<0.001], where the participants continued to have increased expectation for the CS+Vic as compared to the CS- yet trending decreasing expectancy by late trials for each CS type. During extinction recall, participants reported the highest shock expectancy for the first trial of the CS+Vic, followed by the CS+Dir, however stimulus differences disappeared by the last trial [F(2,350)=13.82,*p*<0.001].

**Table 1:**
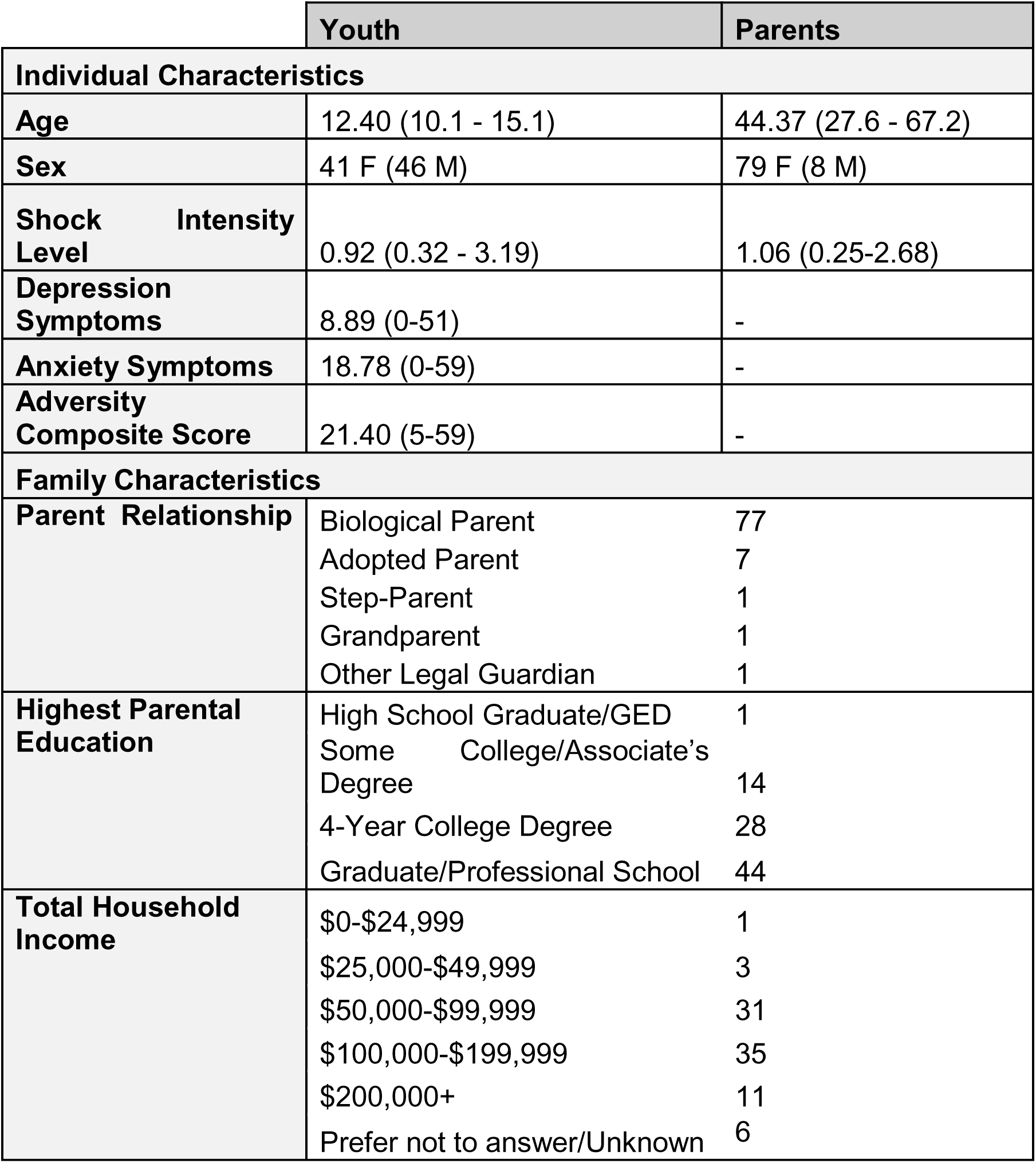
Cohort Sociodemographics Table. Averages and standard deviations are reported, where appropriate. Depression symptoms are total scores from the Mood and Feelings Questionnaire (MFQ). Anxiety symptoms are total scores from the Screen for Child Anxiety Related Disorders (SCARED). The Adversity Composite Score is calculated from the clinician-administered Yale-Vermont Adversity in Childhood Scale (Y-VACS).

### Autonomic Physiology

Mirroring behavioral analyses, SCR provides additional evidence of the successful induction and extinction of threat associations for both CS+ stimuli (Figure 2). A significant stimulus by order interaction during threat acquisition [F(2,913)=6.05,*p*=0.002] detected increased SCR to both the CS+ as compared to the CS-, suggesting increased autonomic arousal and therefore successful threat association, and overall decreasing SCR for all stimuli throughout the phase. While there were no significant stimulus by trial number interactions during direct [F(1,348)=0.0004,*p*=0.98] or vicarious [F(1,312)=0.47,*p*=0.49] extinction, youth exhibited no unique effects of stimulus or trial number in arousal during extinction recall on Day 3 [F(2,892)=0.67,*p*=0.51], suggesting a successful extinction of US-CS associations for both the CS+Vic and CS+D.

**Figure 2.**
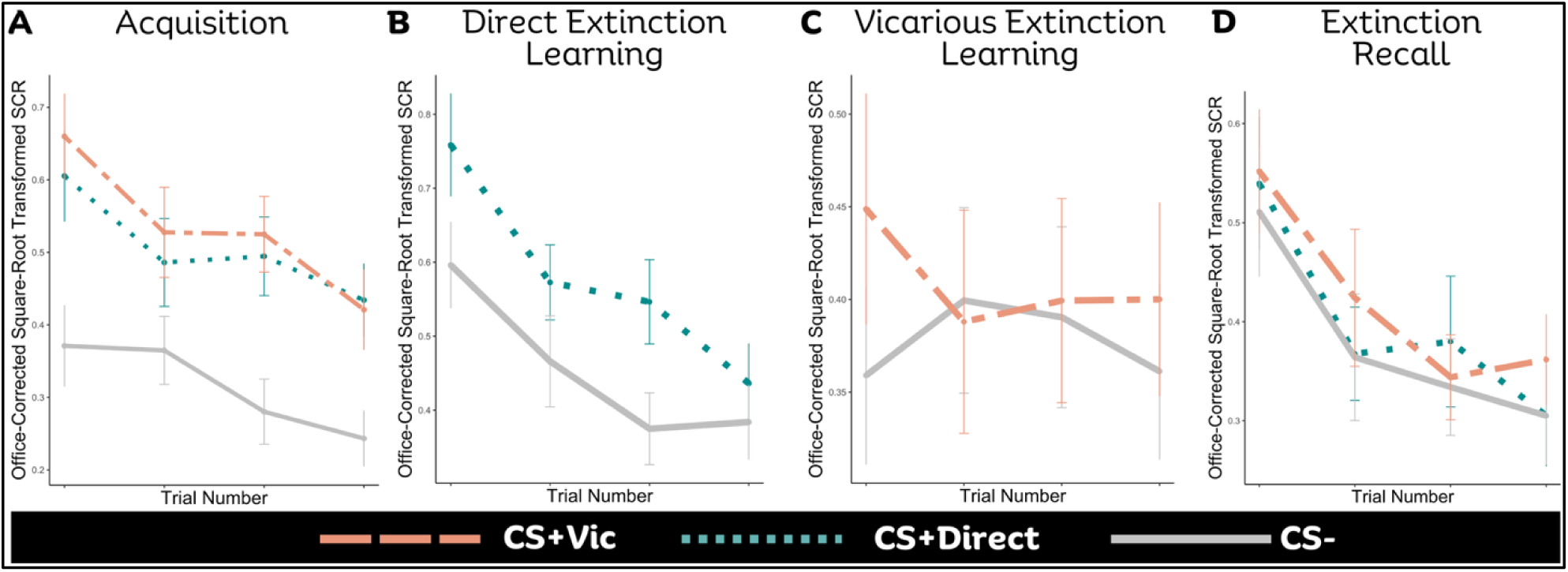
Trajectories of autonomic arousal during the observational threat extinction paradigm. Average SCR response (corrected for the immediately preceding scene and square-root transformed) to each of the first four trials of each stimulus type during acquisition (A), direct extinction (B), vicarious extinction (C), and extinction recall (D). The CS- is depicted by the solid gray line, the CS+Dir is depicted by the dotted blue line, and the CS+Vic is depicted as the dashed orange line. Error bars represent the standard error.

Within parent-child physiological synchrony, the first principal component of the PCA conducted on determinism (DET), laminarity (LAM), and entropy (ENTR) measures explained 90% of the variance. As validation for the component, when investigating whether the synchrony predicts arousal during the vicarious extinction and extinction recall phases, we detected significant relationships with average SCR during the last two trials of both the CS+Vic [F(1,62)=12.47,*p*=0.0008] and CS- [F(1,62)=7.44,*p*=0.008] of vicarious extinction (Figure 4C). Interestingly, greater parent-child synchrony predicted greater autonomic arousal (i.e. greater threat response) to both threat and safety-related cues. Synchrony did not significantly predict SCR for any stimulus during early trials of extinction recall [all *p* values > 0.1].

### Unique patterns of neural activation during an observational threat extinction paradigm

Next, we investigated BOLD activation during each phase of the paradigm while controlling for youth age, sex, cumulative adversity, and PTSD symptoms. Significant results from both whole-brain voxelwise and ROI analyses for each phase are presented in Table 2 and summarized here. First, voxel-wise linear mixed effect modeling using a whole-brain approach examined phase by stimulus type by trial order interactions in BOLD activation. A significant three-way interaction within a diffuse, whole-brain cluster (k=138,753, p_corr_<0.001) was detected, accompanied by widespread significant three-way interactions within 61 of 66 ROIs tests [all *p_corr_*<0.05], spurring targeted decomposition analyses within each phase.

**Table 2.** Significant Stimulus by Order Interactions from Whole-Brain and Region-of-Interest BOLD Activation Analyses. *Abbreviations*: WB, whole brain; ROI, region-of-interest; B, bilateral; L, left; R, right; dACC, dorsal anterior cingulate cortex; dmPFC, dorsomedial prefrontal cortex; vlPFC, ventrolateral prefrontal cortex; vmPFC, ventromedial prefrontal cortex.

| Phase | R/L | Region | WB or Atlas ROI | Cluster Size (k) | x | y | z | p |
| --- | --- | --- | --- | --- | --- | --- | --- | --- |
| Acquisition | B | Whole Brain | WB | 169,854 | 25 | -42 | 6 | <<0.001 |
|  | L | Anterior Hippocampus | Tian 28 | - | - | - | - | <<0.001 |
|  | L | Lateral Amygdala | Tian 48 | - | - | - | - | <<0.001 |
|  | L | dACC | Schaefer 108 | - | - | - | - | <<0.001 |
|  | L | dmPFC | Schaefer 152 | - | - | - | - | <<0.001 |
|  | L | vIPFC | Schaefer 128 | - | - | - | - | <<0.001 |
|  | L | vmPFC | Schaefer 161 | - | - | - | - | <<0.001 |
| Direct Extinction | R | Posterior Insula | WB | 81 | 45 | -5 | 8 | 0.05 |
|  | L | Posterior Insula | WB | 80 | -38 | 3 | 4 | 0.05 |
|  | R | vmPFC | Schaefer 369 | - | - | - | - | 0.03 |
| Vicarious Extinction | R | vIPFC | WB | 97 | 43 | 43 | -2 | 0.04 |
|  | R | vIPFC | Schaefer 305 |  |  |  |  | 0.05 |
| Extinction Recall | - | None | WB |  |  |  |  |  |
|  | L | vmPFC | Schaefer 111 | - | - | - | - | 0.01 |
|  | L | vIPFC | Schaefer 141 | - | - | - | - | 0.03 |
|  | R | Medial Amygdala | Tian 22 | - | - | - | - | 0.04 |
|  | L | vmPFC | Schaefer 109 | - | - | - | - | 0.04 |
| Reinstatement | - | None | WB |  |  |  |  |  |
|  | R | Anterior Hippocampus | Tian 1 | - | - | - | - | 0.03 |
|  | R | dmPFC | Schaefer 377 | - | - | - | - | 0.04 |
|  | R | dACC | Schaefer 373 | - | - | - | - | 0.05 |
|  | R | vmPFC | Schaefer 318 | - | - | - | - | 0.05 |

As expected, no whole-brain clusters or ROIs exhibited stimulus by trial order interactions survived correction during habituation. Threat acquisition, however, again exhibited extensive stimulus by trial interactions in both analyses. Here, a brain-wide cluster [k=169,854, *p_corr_*<0.001] and 63/66 [all *p_corr_*<0.05] ROIs indicate stimulus by trial order effects of BOLD activation, suggesting differential early neural encoding of threat association (Figure 3A). Specifically, across all clusters and ROIs, despite randomization of stimulus presentation in acquisition, regions uniformly inversely activate during the first two trials of each CS+ stimulus (CS+Vic show increased activation as compared to the CS-, while CS+D show decreased activation as compared to the CS-), and these early encoding differences dissipate by late acquisition.

**Figure 3.**
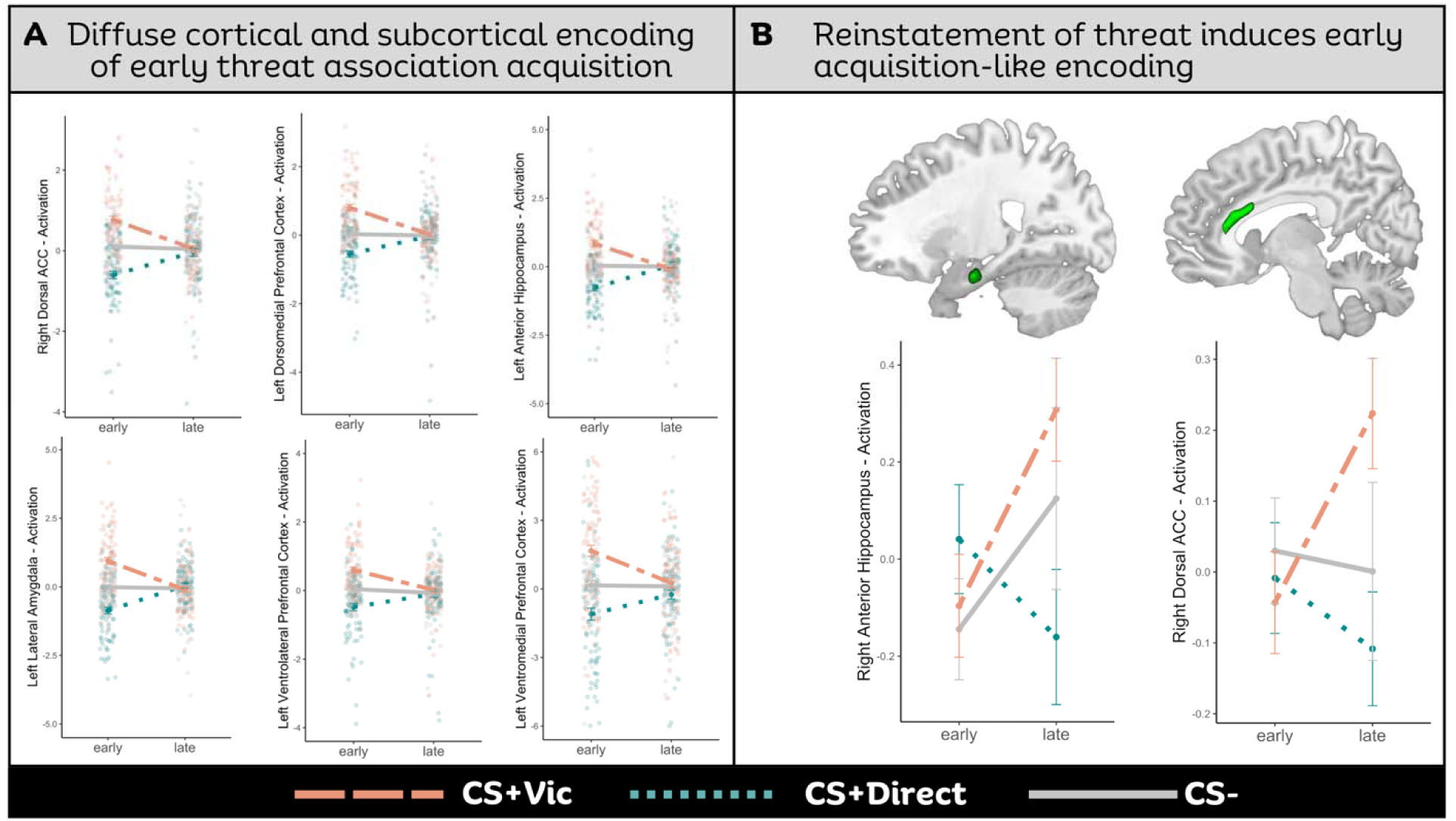
Early differential neural encoding of visual stimuli during threat acquisition mirrors the encoding following reinstatement of threat associations. (A) Average BOLD activation in early versus late phase acquisition to the CS- (solid gray), CS+Dir (dotted blue), and CS+Vic (dashed orange) across numerous brain regions, including (from left to right): the right dorsal anterior cingulate cortex (dACC), left dorsomedial prefrontal cortex (dmPFC), left anterior hippocampus, left lateral amygdala, left ventrolateral PFC (vlPFC), left ventromedial PFC (vmPFC). Data points represent individual activation betas for each participant in early and late phase acquisition overlayed by the corresponding linear line of best fit. (B) The reinstatement of acquisition-induced patterns of BOLD activation in the right hippocampus (left panel) and dACC (right panel) during late reinstatement, where cohort averages and corresponding standard error bars are plotted.

**Figure 4.**
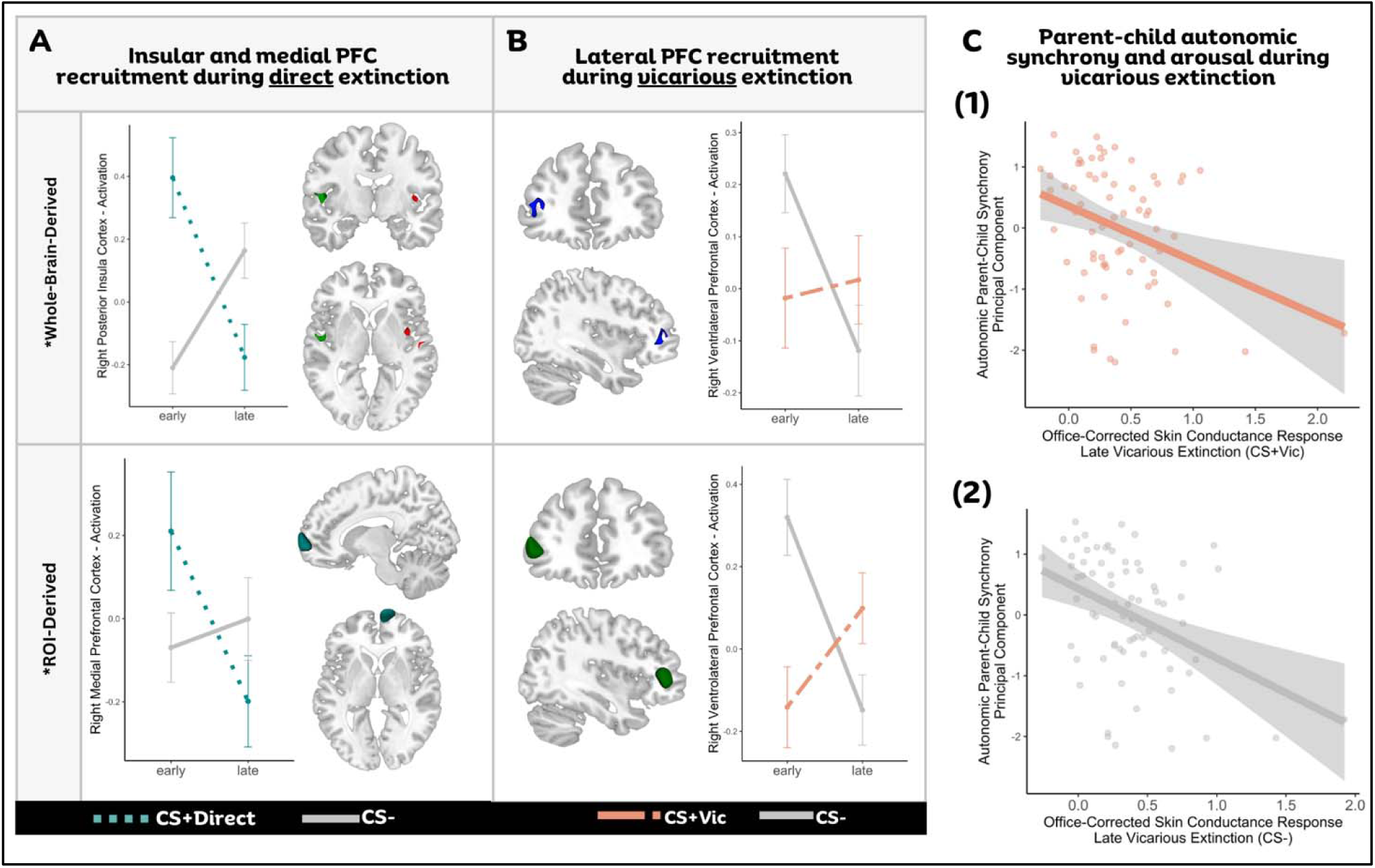
Unique prefrontal recruitment during direct versus observational extinction learning and underlying parent-child autonomic synchrony. (A) BOLD activation patterns, visualized by cohort averages and corresponding standard error bars, during early and late phase direct extinction, controlling for age, sex, and subject as a random effect. The top panel represents significant effects within the bilateral posterior insula detected by whole-brain analyses. The bottom panel represents significant effects within the right ventromedial prefrontal cortex (vmPFC) detected by region-of-interest (ROI) analyses. (B) BOLD activation patterns, visualized by cohort averages and corresponding standard error bars, during early and late phase vicarious extinction, controlling for age, sex, and subject as a random effect. Both top (whole-brain derived) and bottom(ROI-derived) panels depict significant effects within the ventrolateral PFC (vlPFC). (C) Significant negative correlations between the composite of parent-child autonomic synchrony and late-phase vicarious extinction arousal (average SCR in the last two trials of a stimulus, corrected for peak SCR during preceding scene and square-root transformed) during CS+Vic presentation (1) and CS- presentation (2).

During direct extinction, whole-brain analyses detected significant stimulus by order interactions in the bilateral posterior insula [left, k=81, (45, -5, 8), *p_voxelwise_*<0.005, *p*_corr_<0.05); right, k=80, (-38, 3, 4), *p_voxelwise_*<0.005,*p_corr_*< 0.05] (Figure 4A). Here, the posterior insula displays higher recruitment during the early CS+Dir and is decreased across the phase, while the opposite relationship is seen for the CS- trials. ROI analyses additionally detect a stimulus by order interaction in the right ventromedial prefrontal cortex (vmPFC) [F(1,234)=4.93, *p*=0.027] (Figure 4A). Within the vmPFC, recruitment patterns show a similar relationship, where activation to early CS+Dir trials is greater than CS- trials and decreases over time.

During parent-child vicarious extinction training, we see unique effects within the lateral PFC (Figure 4B). Whole-brain analyses detect a stimulus type by trial order interaction in the right ventrolateral PFC (vlPFC) [k=97, (43,43,-2), *p_voxelwise_*<0.005, *p*_corr_<0.04). A corresponding right vlPFC region from ROI analyses reveal an analogous stimulus by trial order interaction [F(1,227)=3.57, *p*=0.05]. In both cases, we see increasing recruitment of the vlPFC to the CS+Vic trials and decreasing recruitment during the CS- trials.

Whole-brain analyses of extinction recall and reinstatement phases on Day 3 did not detect any significant clusters for stimulus type by trial order interactions [all *p_corr_*>0.10]. Among ROI analyses for extinction recall, the left vmPFC [cluster 1, F(2,372)=4.32,*p*=0.01; cluster 2, F(2,372)=3.18,*p*=0.04], left vlPFC [F(2,372)=3.56,*p*=0.03], and right medial amygdala [F(2,373)=3.21,*p*=0.04] all exhibit significant stimulus type by trial order interactions (Figure 5). Within these regions, recruitment during CS- trials increased across extinction recall while recruitment during CS+Vic decreased over time. Interesting, for CS+Vic trials, activation of the amygdala similarly decreased over time, but there was no significant effect during CS+Dir trials, suggesting unique correlates of direct and vicarious extinction recall in the lateral PFC.

**Figure 5.**
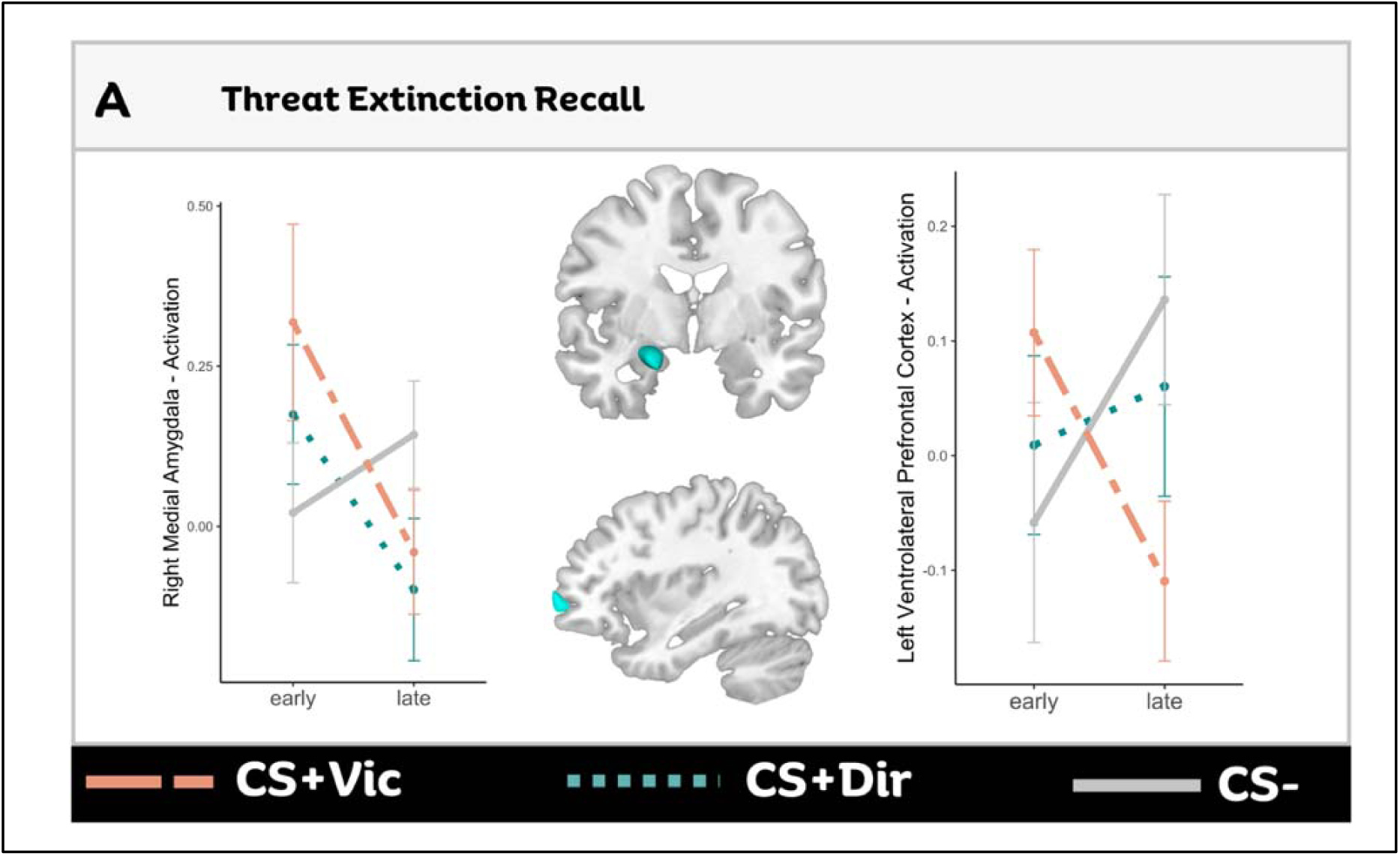
Neural representation of threat extinction recall. (A) BOLD activation patterns, visualized by cohort averages and corresponding standard error bars, during early and late phase extinction recall, controlling for age, sex, and subject as a random effect. The left panel represents a significant effect within the right medial amygdala and the right panel represents significant effect in the ventrolateral prefrontal cortex (vlPFC).

Finally, the reinstatement of threat associations through the delivery of unpaired shocks prior to task start induced unique patterns of activation in hippocampus and medial PFC (Figure 3B). ROI analyses detected significant stimulus type by order interactions in the right anterior hippocampus [F(2,367)=3.50,*p*=0.03], dorsal ACC [F(2,437)=2.99,*p*=0.05], right dmPFC [F(2,368)=3.33,*p*=0.04], and right vmPFC [F(2,437)=2.98,*p*=0.05]. Mirroring the early encoding patterns of threat acquisition, in late trials of reinstatement, each region exhibits differential recruitment of CS-type where the CS+Vic trials exhibit increased activation and the CS+Dir trials exhibit decreased activation as compared to the CS- trials.

## IV. DISCUSSION

In this study, we employ a comprehensive assessment of direct and vicarious threat extinction within youth and their caregivers with the purpose of building a foundation for understanding the mechanisms of the creation and modification of threat associations in youth, regardless of psychopathology and traumatic and adverse early childhood experiences. Although there has been consistent discussion of ethical concerns[45,46] and low retention[47],[48],[49] of youth participating in threat extinction paradigms, we show behavioral, physiological, and neural substrates of threat acquisition and vicarious extinction within parent-child dyads using a highly tolerable paradigm[4] with minimal dropout. Here, we report three key findings. First, associative learning using a highly salient unconditioned stimulus (US; electrodermal stimulation) in youth resulted in robust, early encoding of threat via differential activation patterns across cortical and subcortical regions that is mirrored during threat reinstatement. Second, we provide corroborative evidence of the ventromedial prefrontal cortex (vmPFC) during direct extinction learning in adolescents and suggest expansion of this extinction network to include the ventrolateral PFC (vlPFC) during vicarious extinction processes. Finally, we report one possible mechanism of vicarious extinction learning to be dyadic physiological synchrony, which was negatively predictive of youth arousal to both threat and safety stimuli.

Threat detection and associative learning are stable, highly conserved processes that signal the body to appropriately respond to threatening stimuli[50] and employ the fear extinction network (FEN)[10,12]. In particular, the basolateral amygdala complex is a central hub for converging sensory and contextual information from thalamic and hippocampal inputs[51], while the central nucleus of the amygdala is the locus of threat response initiation[52,53]. Here, projections to the hypothalamus and brainstem to initiate the canonical physiological threat response (e.g., increased heart rate, skin conductance response, freezing behaviors, etc.)[52–54]. As expected, our paradigm successfully induces CS-US associative learning as seen by significantly higher SCR and behavioral shock expectancy to each CS+ type during acquisition in both parents and youth. Neural activation analyses during early acquisition corroborate the recruitment of the amygdala and hippocampus in forming these threat associations, where regions exhibit early differentiation of the CS+Vic and CS+Dir as compared to no distinctive FEN recruitment to the safety cue (CS-). Because CS+ presentation order was randomized, this uniformity of encoding direction likely holds functional significance related to differences in the visual signals. The CS+Vic robustly induced higher activation than safety cues while the CS+Dir elicited lower activation, which may reflect the highly conserved saliency of the color red (CS+Vic), which has been described to uniquely attract attention and reactivity during emotional experiences[55]. Further, this homogenous pattern of activation to CS+ types was detected across many cortical and subcortical regions. As adolescence is inherently a time for heightened learning and reactivity[56], this diffuse response perhaps emphasizes the sheer salience of threat that adaptively triggers an early whole-brain response that rapidly dissipates as youth learn to effectively discern threat and safety cues with repeated associations, regardless of exposure to trauma or PTSS. While this pattern of rapid habituation during threat acquisition has been previously observed in the adult amygdala[57,58], neuroimaging studies of threat acquisition within normative youth populations are less frequent with no clear patterns of encoding, necessitating ongoing investigation of these hypotheses.

The strength and significance of these encoding patterns is further supported by the reproduction of the same patterns following threat reinstatement within the dorsal anterior cingulate cortex and hippocampus, regions acutely involved in the neural circuitry of threat extinction[59]. Reinstatement, or the presentation of unexpected and unpaired US following extinction, is a powerful tool to assess the ongoing vulnerability and generalizability of previously learned threat associations[29]. Here, we see echoes of earlier encoding patterns in both directly and vicariously extinguished CS+ stimuli, suggesting some normative level of generalized extinction fragility in youth. It is unclear from the present data whether this reinstatement of encoding patterns was due to the strength of the original CS–US association, strength of extinction learning, or an overall neutralization of the affective value of the CS+.

The novelty of this paradigm lies within the multimodal extinction phases, where youth completed both direct (repeated stimulus exposure with no shocks) and vicarious extinction (watching a video of their parent complete direct extinction). During the former, we observed increased early recruitment of the right vmPFC and posterior insula to CS+Dir presentation, as compared to CS-, that dissipated over time. The vmPFC plays a dynamic and adaptive role in the active suppression of conditioned responses[59–61], and the relative decrease and lack of differential stimulus effects by late-phase extinction may actually reflect extinction success. The vmPFC is a known responder to safety cues[62] and extinction innately implies the creation of CS+ as a new safety signal[59], altogether suggesting that youth successfully recruit the vmPFC to suppress initial conditioned fear responses in early phases and have a new association of safety with the CS+Dir in late phases. While a recent meta-analysis purports a more cautionary view of the vmPFC (and amygdala) during extinction learning rather than extinction recall and suggest the need to focus on wider network activity rather than individual nodal activation[14], this pattern of preferential recruitment of the vmPFC to the CS+ continues to be well documented in human and animal models of threat extinction[28,61] as a part of the wider FEN. Furthering this hypothesis, to our knowledge, this is one of the first studies within a developmental population to provide direct evidence corroborating the role of the vmPFC in direct extinction learning. Interestingly, adolescent vicarious extinction learning seemed to circumvent the canonical extinction circuitry as we did not detect any involvement of the vmPFC. In contrast, during the intergenerational social transmission of safety, we see significantly increased BOLD response in the ventrolateral PFC (vlPFC) to early trials of the safety cue (CS-) that rapidly diminished and conversely low recruitment to the CS+Vic that increased across trials. It is presumed that underlying vicarious extinction learning success is the ability to attribute the observation of someone’s emotions to a specific cause (i.e., causal emotion attribution, EA). The EA brain network, particularly during casual attribution processes, involves recruitment of the vlPFC, dmPFC, and superior temporal sulcus, among others[63]. Notably, the vlPFC is thought to aid in making causal social judgements[64,65] and causally supports emotion reappraisal[66], which may explain why recruitment of the vlPFC during CS+Vic trials increased across phase as youth continually gather information to successfully attribute parental emotions to new safety associations. This novel characterization of the neural circuitry of vicarious extinction learning in adolescents supports the theory that vicarious extinction is not a simple mirror of direct extinction but rather involves the concerted cortical recruitment of EA networks to make threat and safety judgements.

We and others have previously hypothesized that the degree of physiological synchrony between the demonstrator and observer, or the similarity in SCR during the learning phase,[22,33] may explain individual differences in vicarious extinction learning success[4]. Here, we provide additional corroborative evidence of this learning mechanism, where increased synchrony was associated with lower arousal to both threat and safety cues *during* vicarious extinction. In other words, a youth’s ability to “sync up” with their caregiver during vicarious extinction correlated with contemporaneous dampening of their threat response. Interestingly, synchrony was not predictive of arousal during extinction recall, perhaps suggesting a generalization of safety across all stimuli by the third day or the presence of unknown but complementary mechanisms for predicting extinction lability or sucess. Others have suggested that the significance of synchrony is highly dependent upon the family context, such as attachment or parenting styles[22,67], which was unexplored here. Alternatively, perhaps the strength of dyadic synchrony and success of vicarious extinction learning also rely upon one’s ability to identify emotions regardless of the inter-dyadic parental relationship. In adolescents, vicarious threat acquisition has been shown to be similar in strength with parent or stranger demonstrators[5], has previously been linked to gaze patterns,[20] and atypical emotion recognition has been reported in youth with disordered threat responses and PTSD[68], but these interindividual factors have yet to be directly investigated. Altogether, we hypothesize that a youth’s ability to match their parent’s biological state may, in part, facilitate the social adaptation of threat and represents an intervenable relationship that is already often targeted in therapeutic interventions[69].

The findings of this study should be viewed considering the following limitations. Despite statistical controls, this was not a true community sample as recruitment for this study was enriched for trauma exposure and/or PTSD and therefore may not be reflective of normative community prevalence rates. While this may limit generalizability, this cohort provides a unique ability to control for patterns associated to trauma and/or PTSD that may be masked within a true community sample and elucidate the robust, common patterns of vicarious extinction in adolescence. Further work directly targeting the impacts of psychopathology and early childhood experiences on vicarious extinction processes within this sample is warranted. Next, the relatively narrow age range of youth within this study (10-14) obscures our ability to identify broader developmental trajectories of extinction processes. While direct learning may be stable between young children and adolescents[70], many others have suggested age-related changes in extinction success between early and late adolescence[71] and known variations in the parent-child relationship[72,73], therefore justifying continued investigation of vicarious extinction and age. Next, this study investigated only voxel- and ROI-level activation and resulting interpretations regarding the threat and emotion circuitry are based solely upon *a priori* hypotheses. Functional connectivity and causal modeling using advanced statistical approaches are needed to characterize the role of the FEN, EAN, and related networks in parent-child vicarious extinction. Given known differences in threat and safety discrimination[74] and prevalence rates of PTSD[75] between male and female youth, work should investigate (rather than control for) differences in biological sex. Finally, there is burgeoning discussion of the ecological validity of simplistic, Pavlovian conditioning using static visual stimuli limits the generalizability of these processes to real-life experiences. Future work expanding these paradigms using ecological momentary assessment (EMA) in emotional reactivity[76] and neurobiology to threatening stimuli during real-life or virtual-reality paradigms may boost generalizability.

To our knowledge, this study is one of the first to examine normative patterns of direct and vicarious extinction in a community sample of adolescents, controlling for previous traumatic and adverse experiences and PTSS severity. We affirm the ability of youth to modify previously-learned threat associations either directly or by the social transmission of safety via parental modeling, detect widespread early encoding of threat, provide corroborative evidence of the role of the vmPFC during direct extinction[10,17] and suggest recruitment of the lateral PFC as critical for vicarious extinction, and propose a possible mechanism of vicarious extinction in parent-child dyadic synchrony. These findings provide the necessary foundation for understanding how trauma and fear-related disorders may ensue with disruption of these processes. Furthermore, potential neurophysiological markers of atypical vicarious extinction may allow for objective, rational targeting of treatment in the parent-child dyad to improve outcomes within these vulnerable populations.

## ACKNOWLEDGEMENTS

We owe our sincerest gratitude to the youth and families who have given their time for this study. Funding for this study was provided by the National Institutes of Health/National Institutes of Mental Health (K08MH100267, R01 MH117141, R01 MH115910, to RJH), American Academy of Child and Adolescent Psychiatry Junior Investigator Award (to RJH), NARSAD Young Investigator Grant (to RJH), University of Wisconsin Institute for Clinical and Translational Research Translational Pilot Grant Award (NIH/NCATS UL1TR000427, to RJH), University of Wisconsin Institute of Clinical and Translational TL1 Training Award(TL1TR000429, to SAH), and the University of Wisconsin School of Medicine and Public Health. None of these funding sources had a direct effect in the design, analysis, or interpretation of the study results, nor in preparation of this manuscript.

## DATA AVAILABILITY

This data was collected under R01-MH117141. As such, much of the participant demographic, clinical, behavioral, and neuroimaging data will be uploaded and available on the National Institute of Mental Health Data Archive (NDA). Additional data available upon reasonable request to the corresponding author (Heyn).

